# Human CDK5–cyclin B1 structures uncover a conserved mitotic CDK activation mechanism

**DOI:** 10.1101/2025.09.10.675329

**Authors:** Andrew S. Arvai, Ke Cong, Xiao-Feng Zheng, Amy Verway-Cohen, Miaw-Sheue Tsai, Huy Nguyen, Albino Bacolla, Chi-Lin Tsai, George Lantz, Alexander Spektor, Dipanjan Chowdhury, John A. Tainer, Aleem Syed

## Abstract

Cyclin-dependent kinase 5 (CDK5), long considered atypical and activated by non-cyclin cofactors in neurons, also functions in mitosis. To resolve its activation mechanism during mitosis, we determined the first high-resolution crystal structures of CDK5–cyclin B1 in apo and nucleotide-bound states. Contrary to AlphaFold predictions, we find CDK5 structures unexpectedly mirrors CDK1-cyclin B1 assembly and activation, establishing CDK5 as a *bona fide* mitotic kinase acting in parallel with CDK1.

---

The cell cycle is a precisely regulated process that ensures accurate replication and division of cells, resulting in two identical daughter cells^1^. Central to this regulation are cyclins— nonenzymatic proteins whose abundance oscillates across the cell cycle phases^2^. Cyclins activate CDKs, which are proline-directed serine/threonine kinases essential for cell cycle progression^3^. Of the many CDKs and cyclins identified, only a subset directly regulates cell cycle transitions. CDK2, CDK3, CDK4, and CDK6 control interphase progression, whereas CDK1 is essential for mitosis^4^. Uniquely, CDK1 binds all cell cycle-specific cyclins, including cyclin B1, making it indispensable for mitotic entry and completion^5^. In contrast, the hallmark of atypical CDK5, despite its sequence similarity to CDK1 and CDK2 (Extended Data Fig. 1), is its activation by non-cyclin proteins p35/p25 and p39/p29 in post-mitotic neurons, primarily influencing neuronal development^6,7^.

We recently uncovered crucial roles for CDK5 in mitosis^8,9^, where its inhibition or depletion compromises mitotic fidelity^9^. During mitosis, CDK5 associates with cyclin B1, a canonical activator of CDK1. However, structural predictions have been inconsistent with this role: AlphaFold2 (AF2) modeling suggested that the CDK5–cyclin B1 complex more closely resembles CDK2–cyclin A2^9,10^. Moreover, AlphaFold3 (AF3) predicted a structure of the CDK5-cyclin B1 identical to the AF2 model with a low-confidence addition to the activation loop in CDK5 and to the extreme cyclin N-terminus (Extended Data Fig. 2). Here, we report the first experimental crystal structures of the CDK5–cyclin B1 complex. Our data reveal that the active assembly mirrors the canonical CDK1–cyclin B1 complex, thereby establishing the molecular mechanism by which CDK5 functions as a *bona fide* mitotic kinase.

Our biochemical analyses (reciprocal pull-down assays from two different insect cell lines, gel-filtration chromatography, and kinase activity assays) found that CDK5 and cyclin B1 form an active stoichiometric complex when co-expressed and purified under near-physiological salt conditions (Fig. 1a,b and Extended Data Fig. 3). Accordingly, we crystallized the recombinantly purified complex and solved CDK5-cyclin B1 structures with and without bound AMP-PNP, a non-hydrolyzable ATP analog (Fig. 1c-g, Extended Fig. 4, and Table 1). Corroborating the biochemical data, the crystallographic structures reveal that CDK5 adopts an active conformation when complexed with cyclin B1. Key features indicated CDK5 activation: an extended activation loop (A-loop, D144-P170), alignment of regulatory spine residues (L55, L66, H124, and F145) due to inward C-helix displacement upon cyclin B1 binding (P45-E57, nomenclature from first protein kinase structure^11^), and characteristic tripeptide motif (D144-F145-G146) in the “DFG-in” configuration for active kinases^12–14^ (Fig. 1c-e).

**Table 1.** X-ray data collection and refinement statistics for CDK5-cyclin B1 structures. Statistics values for the highest resolution shell are in parentheses.

|  | CDK5-cyclin B1<br>(apo) | CDK5-cyclin B1<br>(AMP-PNP) |
| --- | --- | --- |
| **Data Collection Statistics** |  |  |
| Wavelength (Å) | 0.9795 | 0.9795 |
| Resolution range (Å) | 38.98 - 2.10 (2.16 - 2.10) | 39.56 - 2.5 (2.6 - 2.5) |
| Space group | P 32 2 1 | P 32 2 1 |
| a, b, c (Å) | 121.58 121.58 115.99 | 120.86 120.86 116.07 |
| $\alpha, \beta, \gamma$ (°) | 90 90 120 | 90 90 120 |
| Total reflections | 592857 (45806) | 355840 (38497) |
| Unique reflections | 58032 (4423) | 34310 (3800) |
| Multiplicity | 10.2 (10.4) | 10.4 (10.1) |
| Completeness (%) | 99.8 (99.2) | 99.9 (99.6) |
| Mean I/sigma(I) | 13.1 (0.4) | 9.4 (0.5) |
| Wilson B-factor | 51.77 | 50.59 |
| R-merge | 0.078 (7.133) | 0.215 (4.712) |
| CC1/2 | 0.999 (0.281) | 0.998 (0.287) |
| **Refinement Statistics** |  |  |
| Reflections used in refinement | 47468 | 28950 |
| Reflections used for R-free | 1658 | 1441 |
| R-work | 0.17 | 0.19 |
| R-free | 0.22 | 0.25 |
| Number of non-hydrogen atoms | 4766 | 4710 |
| Protein | 4503 | 4515 |
| Water | 164 | 71 |
| Other | 99 | 124 |
| RMS (bond length) (Å) | 0.004 | 0.002 |
| RMS (bond angle) (°) | 0.610 | 0.528 |
| Ramachandran favored (%) | 94.95 | 95.68 |
| Ramachandran allowed (%) | 5.05 | 4.32 |
| Ramachandran outliers (%) | 0 | 0 |
| Rotamer outliers (%) | 3.82 | 5.81 |
| Clashscore | 4.31 | 5.35 |
| Average B-factor | 95.42 | 76.69 |
| Protein | 94.91 | 76.25 |
| Water | 88.01 | 63.89 |
| Other | 131.00 | 100.13 |
| PDB | 9Q3N | 9Q3O |

**Fig. 1.**
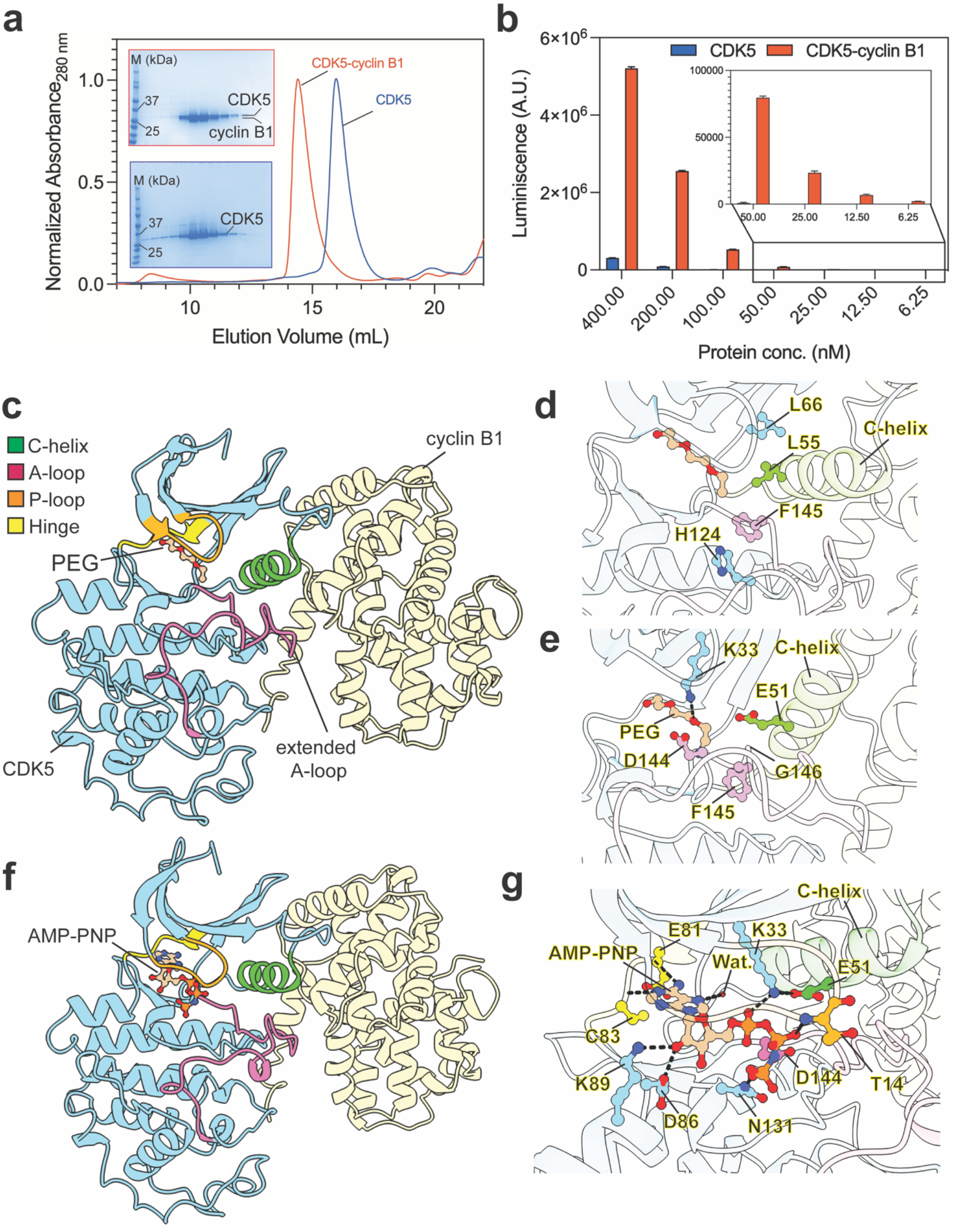
CDK5 unexpectedly forms an active canonical CDK-cyclin complex structure with cyclin B1. a) Gel-filtration profiles show CDK5 elutes as a stable heterodimer with cyclin B1 but as a monomer alone. Protein gels are shown for the peak fractions (fraction# 23-34 for CDK5-cyclin B1 or 28-38 for CDK5 only). b) Kinase activity at varying concentrations of the CDK5-cyclin B1 complex confirms CDK5 activation by cyclin B1. c) The CDK5-cyclin B1 crystal structure reveals an active conformation of CDK5 with an extended activation loop, and the C-helix-in configuration. PEG occupies the ATP binding site. d) Alignment of regulatory spine residues (RS1-H124 C-ter lobe, RS2-F145 from DFG motif, RS3-L55 from C-helix and RS4-L66 from N-ter lobe) in the active complex. e) The characteristic salt bridge between K33 (from the β2-strand) and E51 (from the C-helix) is disrupted by PEG hydrogen bonds with K33. f) The AMP-PNP-bound CDK5-cyclin B1 structure shows ATP binding in the active site. g) Interaction network formed between AMP-PNP and CDK5 in the active site. The critical regions (C-helix, A-loop, P-loop and hinge) are labeled according to the color legend (shown in panel c) in the structures.

In the apo structure, a poly(ethylene glycol) (PEG) molecule occupies the active site, forming a hydrogen bond with catalytic lysine residue K33, thereby disrupting the canonical salt bridge between K33 and E51 within the CDK5-cyclin B1 complex (Fig. 1e). Conversely, in the AMP-PNP-bound configuration, the K33-E51 salt bridge is reestablished and further engages with the phosphate moiety of the bound ligand within the active site, consistent with active kinase conformations (Fig. 1g). The AMP-PNP-bound structure also elucidates the mode of ATP recognition within the active site of the CDK5-cyclin B1 complex. The adenine moiety of AMP-PNP participates in the classic hinge interactions, while added contacts formed between the CDK5 active site and the ribose ring plus the phosphate groups of AMP-PNP (Fig. 1g). Furthermore, bound AMP-PNP completes the catalytic spine interaction network, signifying the catalysis-ready state of the active CDK-cyclin complex^14^ (Extended Data Fig. 5).

The AF2-predicted model of the CDK5-cyclin B1 complex indicated that the tip of the activation loop, specifically the I153 residue, is stabilized via hydrophobic interactions, resembling those in the CDK5-p25 complex^15^ or the V154 residue in the CDK2-cyclin A2 complex structures^10,16^. However, these residues point in opposite directions due to distinct stabilization mechanisms by p25 or cyclin A2^9^. Contrarily, our crystallographic data reveal that the tip of the activation loop is constituted by the F151 residue, which is stabilized by many productive interactions with cyclin B1 (Fig. 2a,b).

**Fig. 2.**
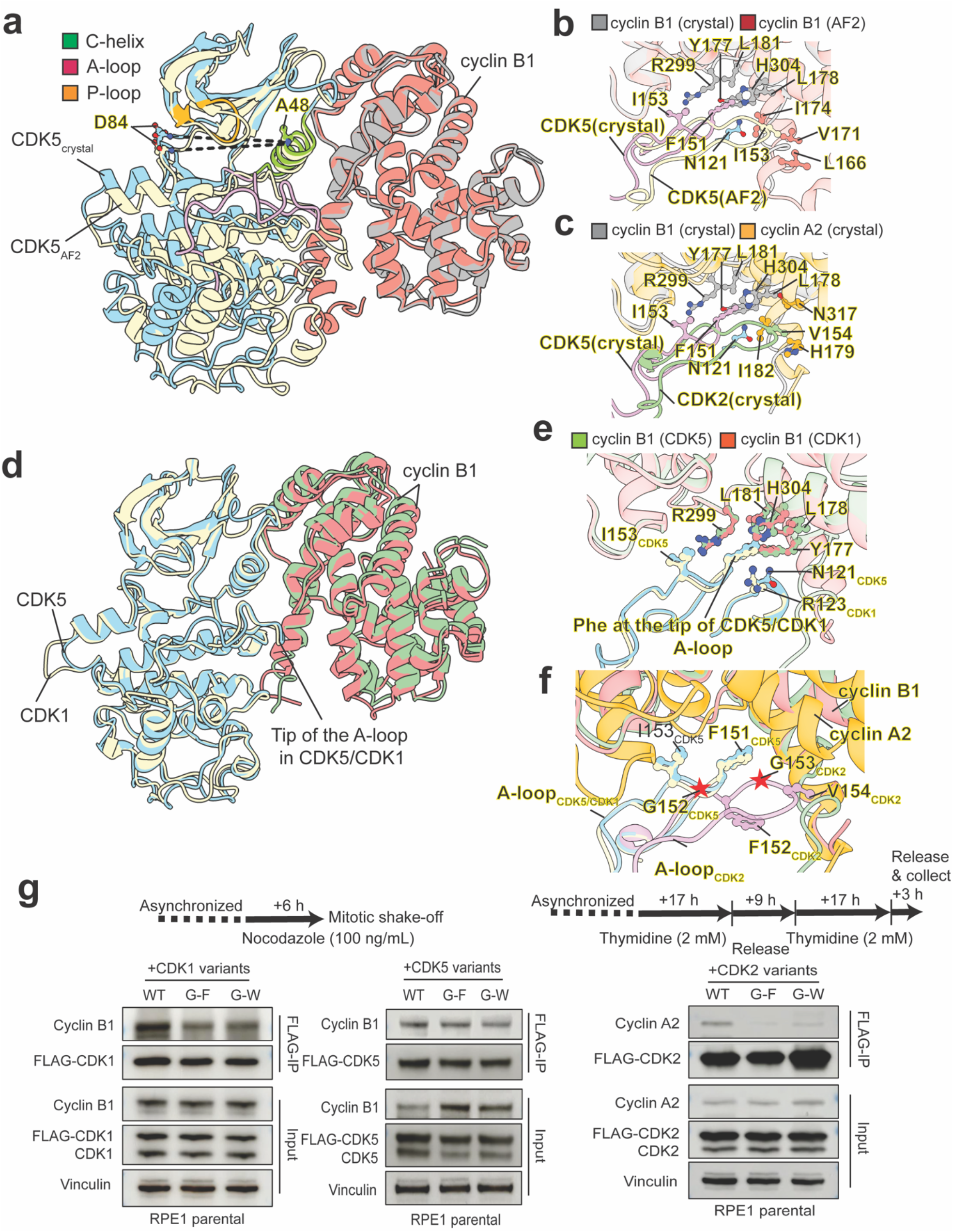
Strikingly conserved assembly and activation mechanisms for CDK5-cyclin B1 structure and assembly with mitotic CDK-cyclin B1 complexes. a) Superposition of the CDK5-cyclin B1 crystal structure with the AF2 model (using cyclin B1 as reference) reveals differences in CDK5 subunit disposition, A-loop stabilization, and active site configuration (distance between A48 and D84 is 22 Å in AF2 and 24 Å in the crystal structure). b) Predictive discrepencies in the AF2 model and stabilization differences in the CDK5 A-loop by cyclin B1: F151 at the tip of the A-loop in the crystal structure (pink) vs I153 at the tip of the A-loop in the AF2 model (wheat). c) Comparison of A-loop stabilization in CDK5 versus CDK2 (PDB: 1QMZ) by cyclin B1 and cyclin A2, respectively. d) Superposition of crystal structures of CDK5-cyclin B1 and CDK1-cyclin B1 (PDB-4YC3) shows conserved assembly. e) Zoomed view of identical A-loop stabilization in CDK5 and CDK1 by cyclin B1. f) Glycine (G152_CDK5_, G154_CDK1_ or G153_CDK2_ residue position (red star) between Phe and Ile/Val in the A-loop of CDK1, CDK2, and CDK5, mutated to bulky residues for studies. g) Synchronization schematic (upper panel) and immunoblots of Flag immunoprecipitation of the indicated Flag-CDK variants expressed in RPE-1, synchronized (to M-phase for CDK1 or CDK5 and S-phase for CDK2) as shown, and blotted for the indicated proteins (lower panel).

Furthermore, superimposition of the crystal structures of CDK5-cyclin B1 with either the AF2-predicted model of CDK5-cyclin B1^9^ or the crystal structure of the CDK2-cyclin A2 complex^10^, using cyclin as a reference, reveals an altered spatial disposition of the cyclin-CDK subunits including differences in the A-loop (Fig. 2b,c and Extended Data Fig. 6a). While cyclin and C-helix exhibit perfect alignment (r.m.s.d. 0.345 Å over 206 equivalent Cα), the entire CDK domain is displaced. Notably, the active site configuration, particularly the distance between the C-helix (A48) and the hinge region (D84), is expanded (24.4 Å) versus the AF2 model or CDK2-cyclin A2 structure (22.2 Å, Fig. 2a and Extended Data Fig. 6a). For alignment by the CDK domain reference, the cyclin failed to align, yet most of the CDK domain aligned (r.m.s.d. 0.508 Å over 222 equivalent Cα), excepting an increased C-helix to hinge region distance within the active site (Extended Data Fig. 6b,c). Significantly, ATP-competitive inhibitors predominantly target this region, so the structural differences inform rational design of CDK-selective inhibitors.

Aligning our structure with the published CDK1-cyclin B1 complex^17^, results in a congruent alignment of both cyclin and CDK domains (r.m.s.d. 0.711 Å over 458 equivalent Cα), including the extended configuration of the A-loop with F153 ( analogous to F151 in CDK5) at the tip (Fig. 2d,e). Interestingly, V154 at the tip of human CDK2 A-loop is aligned to I151 in human CDK5 or I153 in human CDK1, which may have contributed to how A-loop is extended in these kinases, although valine occurs at this Cdk1 position in some species (Extended Data Fig. 7, Supplementary data 1). Resembling the CDK1-cyclin B1 structure, the A-loop residues proximal to the substrate binding regions (C157-L165) exhibit increased lability (average B-factor 149 Å^2^) compared to the remaining activation loop (D144-R156 and W166-P170, average B-factor 78 Å^2^), suggesting flexibility for substrate binding^17^. Notably, the structural conservation between cyclin B1 structures of CDK1 and CDK5 implies CDK5-cyclin B1 complex may also share regulation via A-loop and glycine-rich P-loop phosphorylation as these sites are highly conserved in CDK5 across species, analogous to CDK1 (Extended Data Figs. 7a,b and 8a-d). Supporting this hypothesis, we found that CDK5 P-loop phosphorylation at T14/Y15 diminished as cells enter into mitosis (Extended Data Fig. 9) and our prior results found that the A-loop phosphorylation site (S159) is critical for CDK5-cyclin B1 mitotic function^9^. Collectively, our structural and biochemical data support conserved assembly and activation mechanisms between CDK1 and CDK5 as critical for maintaining mitotic fidelity.

To validate the observed structural differences between mitotic and S-phase cyclin-CDK complexes, we examined CDK5 G152 (corresponding to CDK2 G153 and CDK1 G154), which is positioned away from the cyclin interface in cyclin B1-bound CDK5 or CDK1 structures but oriented towards the cyclin interface in CDK2-cyclin A2 complex (Fig. 2f). We introduced mutations at the G152 residue to bulkier residues (Y or W) across all three kinases. As predicted in RPE1 cells synchronized to their respective cell cycle phases, the bulky mutations at this glycine residue led to a disruption of CDK2-cyclin A2 binding during the S-phase, while showing minimal impact on CDK5/CDK1 binding to cyclin B1 during the M-phase (Fig. 2g).

For structural molecular biology, our findings demonstrate that CDK5, long considered atypical, in fact functions as a canonical mitotic CDK through direct activation by cyclin B1. The crystal structures reveal that the fundamental principles of cyclin–CDK assembly and activation are conserved between CDK1 and CDK5, overturning prior assumptions and diverging sharply from computational predictions. Although AlphaFold produced high-confidence models, our experimental data expose critical inaccuracies in activation loop conformation and interfacial contacts. These discrepancies underscore the indispensable role of empirical validation in the era of AI-based modeling. Notably, AF2 may yield low-confidence models for sequences lacking robust multiple sequence alignment (MSA) or sufficient homologous structural data in the PDB^18–20^. Yet here high-quality prediction was evidently skewed by the disproportionate abundance of structural data for CDK2 in the PDB (>450 structures) relative to CDK1 and CDK5 (~10 structures each), leading the model to resemble CDK2 rather than accurately reflecting the novel CDK5 complex.

For cell biology, our results redefine CDK5 as a *bona fide* mitotic kinase, operating in parallel with CDK1 to safeguard chromosome segregation and mitotic fidelity. By revealing its structural and mechanistic analogy to CDK1, our study expands CDK5’s role in cell cycle regulation and raises the possibility that CDK5–cyclin B1 can supplement for CDK1–cyclin B1 activity under specific conditions, including in cancer and other pathologies characterized by high proliferative demand. Beyond advancing fundamental knowledge, defining the analogous molecular assembly and activation mechanisms of the mitotic CDK5 complex opens new avenues for therapeutic strategies targeting CDK5 in cancer and neurodegenerative diseases. Together, these insights transform the view of CDK5 from atypical outlier to canonical regulator of mitosis.

## Acknowledgements

Authors would like to thank Dr. Benjamin M. Stinson (Dana-Farber Cancer Institute) for providing helpful feedback on the manuscript. A.Syed acknowledges support from the Claudia Adams Barr Program for Innovative Basic Cancer Research and the JCRT Foundation grant. J.A.T. was supported by National Institutes of Health grants (P01 CA092584, R35 CA2204300, and a Robert Welch Chemistry Chair (G-0010). D.C is supported by R01 CA208244 and R01 CA264900, Gray Foundation Team Science Award, DOD Ovarian Cancer Award W81XWH-15-0564/OC140632, Tina’s Wish Foundation, V Foundation Award, and the Claudia Adams Barr Program in Innovation Basic Cancer Research. K.C. was supported by the Postdoctoral Fellowship (PF-23-1145928-01-DMC) from the American Cancer Society.

## Data availability

All crystallographic data and respective structural models presented have been deposited to the Protein Data Bank (PDB) with accession numbers: CDK5-cyclin B1 (9Q3N), and CDK5-cyclin B1-AMP:PNP 9Q3O). Computational models generated during the study have been deposited to ModelArchive repository (modelarchive.org) with access codes: CDK5-cyclin B1 (https://www.modelarchive.org/doi/10.5452/ma-5qxe6, access code: 8vmAKg5Ayt), CDK5 (pT14)-cyclin B1-ATP (https://www.modelarchive.org/doi/10.5452/ma-akm64, ZXFEndVD8Y), CDK5 (pY15)-cyclin B1-ATP (https://www.modelarchive.org/doi/10.5452/ma-g11ot, access code: WThMIdjgPp), CDK5 (pT14/pY15)-cyclin B1-ATP (https://www.modelarchive.org/doi/10.5452/ma-vaaso, access code: B6ireFPD6T), and CDK5 (pS159)-cyclin B1-ATP (https://www.modelarchive.org/doi/10.5452/ma-agke4, access code: 0VCSagJKLX). All other data generated during the study are included in this article or its supplementary information files.

## Methods & Protocols

### Protein expression and purification

The insect cell expression constructs for CDK5 and cyclin B1 were designed based on the AF2-predicted model^1^. The coding regions for CDK5 (full-length) and cyclin B1 (amino acids 152-433, with mutations C167S, C238S, C350S) were codon-optimized and cloned into 438-series Macrobac baculovirus expression vectors^2^: 438-B with a His-tag and 438-C with a His-MBP-N10-tag.

For the reciprocal pull-down experiments, CDK5 and cyclin B1 were co-expressed by co-infecting High-five (HiFi) cells for 45 hours or Sf9 cells for 70 hours with two individual baculoviruses before harvesting. For protein purification, cell pellets were resuspended in lysis buffer (50 mM Tris-HCl, pH 7.5, 150 mM NaCl, 0.3% NP40, 10% glycerol, 1 mM PMSF, 1 mM beta-mercaptoethanol) and supplemented with Roche cComplete, EDTA-free protease inhibitor tablets (Sigma, 11836170001). Cell lysate was prepared using a Dounce homogenizer, followed by brief sonication and clarified by centrifugation. The clarified lysate was incubated with amylose resin (New England Biolabs, E8021L) for 90 minutes on a rotator in a cold room for binding. The resin was subsequently washed, and the bound protein complex was eluted twice in elution buffer (50 mM Tris, pH 7.5, 150 mM NaCl, 0.3% NP40, 10% glycerol, 1 mM beta-mercaptoethanol, and 10 mM maltose), each with a 5-minute incubation on ice. Both eluted fractions and the leftover resins were analyzed by SDS-PAGE with SimplyBlue SafeStain solution (Thermo Fisher Scientific, LC6060).

For CDK5-cyclin B1 expression and purification, both proteins were expressed from a single 438-B vector in Sf9 cells by infecting them with the corresponding baculovirus for 70 hours before harvesting. For protein purification, cell pellets were resuspended in lysis buffer (50 mM Tris, pH 8.0, 500 mM NaCl, 5% glycerol, 1 mM TCEP, and 10 mM imidazole) supplemented with Pierce EDTA-free protease inhibitor tablets (Thermo Fisher Scientific, A32965). Cell lysate was prepared by sonication and clarified by ultracentrifugation. The clarified lysate was incubated with TALON® metal affinity resin (Takara Bio, 635502) for 90 minutes in a cold room for binding. The resin was subsequently washed, and the bound protein complex was eluted in elution buffer (25 mM Tris, pH 8.0, 300 mM NaCl, 2.5% glycerol, 1 mM TCEP, and 300 mM imidazole). The His-tag from the eluted protein complex was cleaved by TEV protease and removed by passing the mixture through the TALON® resin. Finally, the protein complex was loaded onto a pre-equilibrated size exclusion chromatography (SEC) column, Superdex 200 Increase 10/300 GL, in SEC buffer (50 mM Tris, pH 8.0, 200 mM NaCl, 1 mM TCEP), and fractions containing the CDK5-cyclin B1 complex were concentrated and flash-frozen in liquid nitrogen. CDK5 alone was also expressed and purified using the same method as for CDK5-cyclin B1. Purified proteins were analyzed by SDS-PAGE with Coomassie Brilliant Blue G-250 staining.

### Crystallization and data collection

For crystallization experiments, purified CDK5-cyclin B1 was concentrated to approximately 15 mg/mL in SEC buffer. Apo crystals of the CDK5-cyclin B1 complex were grown by vapor diffusion at 20 ºC from a reservoir solution containing 10% PEG 3350, 200 mM imidazole/malate (pH 7.8), and 5% saturated magnesium formate. For cryoprotection, 25% glycerol was added to the mother liquor. AMP-PNP-bound crystals of the CDK5-cyclin B1 complex were grown by vapor diffusion at 20 ºC from a reservoir solution containing 10% PEG 3350, 200 mM imidazole/malate (pH 6.0), and 5% saturated magnesium formate, with the addition of 2.3 mM AMP-PNP to the protein prior to setting the crystallization drops. AMP-PNP-bound crystals were cryoprotected using 33% ethylene glycol in the mother liquor. X-ray diffraction data were collected from a single crystal at beamline 12-2 of the Stanford Synchrotron Radiation Lighsource (SSRL).

### Structure determination, refinement, and analysis

The diffraction data for the apo CDK5-cyclin B1 complex were processed using XDS^3^. The AlphaFold2-predicted model^1^ was used to phase the complex structure by molecular replacement with Phaser^4^, as implemented in the Phenix suite^5^. Structure refinement was performed in phenix.refine^6^, with iterative manual fitting using Coot^7^. To solve the AMP-PNP-bound structure, molecular replacement was performed using the refined apo structure. The diffraction data were processed using XDS, and molecular replacement was conducted with Phaser in the Phenix suite. Structure refinement was done in phenix.refine, with iterative manual model building and fitting in Coot. Ligand restraints for AMP-PNP were generated using phenix.ready_set. Simulated annealing composite omit maps were generated using Phenix suite. All structural analyses were performed with Coot, PyMol (Schrödinger Inc.), or ChimeraX (UCSF)^8^ and figures were generated with either PyMol (Schrödinger Inc.) or ChimeraX (UCSF). Data collection and refinement statistics are summarized in Table 1.

### Multiple sequence alignment and sequence logos

Sequences corresponding to CDK1, CDK2, or CDK5 from the different multicellular eukaryotes groups (Actinopterygii (taxid:7898); Afrotheria (taxid:311790); Amphibia (taxid:8292); Cephalochordata (taxid:7735); Chondrichthyes (taxid:7777); Coelacanthimorpha (taxid:118072); Crocodylia (taxid: 1294634); Echinodermata (taxid:7586); Euarchontoglires (taxid:314146); Laurasiatheria (taxid:314145); Lepidosauria (taxid:8504); Metatheria (taxid:9263); Neognathae (taxid:8825); Palaeognathae (taxid:8783); Passeriformes (taxid:9126); Psittaciformes (taxid:9223); Protostomia (taxid:33317); Testudines (taxid:8459); Xenarthra (taxid:9348)) were obtained from the UniProt database^9^. All the sequences from each group that identified as CDK1 (577 in total), CDK2 (269 in total) or CDK5 (623 in total) were downloaded as three separate sets and corrected for any duplicate sequences. To be comparable with human CDK sequences in length, from each set of sequences of CDK1, CDK2 or CDK5, multiple sequence alignment was iteratively performed only on sequences with maximum length of 320 amino acids (from 542 CDK1 sequences, 235 CDK2 sequences and 533 CDK5 sequences) using the msa Bioconductor R package (version 1.40.0)^10^. In each iteration, a small number of sequences that introduced large gaps in the previous iteration were removed. The consensus sequence was then computed from the msa result (from 494 CDK1 sequences, 214 CDK2 sequences and 405 CDK5 sequences) using the msaConsensusSequence function. The ggseqlogo R package (version 0.2)^11^ was used to plot the sequence logos of the consensus sequences, with the input consensus matrices computed from msa results.

### AlphaFold3 modelling

Protein sequences for CDK5 (accession# Q00535) and cyclin B1 (accession# P14635) were obtained from UniProt database. The protein length for cyclin B1 (152-433 aa) used for AF3 prediction was based on previously identified interaction surface between CDK5 and cyclin B1 using AF2 multimer to avoid non-interacting unstructured N-terminus for clarity. The full-length protein sequence was used for CDK5. The input sequences for CDK5 and cyclin B1 were entered into AF3 server^12^ (https://alphafoldserver.com/) with appropriate post translational modifications (PTMs) on CDK5 and ATP used as a ligand for the structure prediction with default input parameters. For the apo CDK5-cyclin B1 model, PTMs and ATP ligand were omitted from the input. The local confidence in the prediction were indicated by pLDDT score for each amino acid in the output .cif file in the B-factor column and an overall ipTM value. For each prediction, the top model was used for downstream comparative structural analysis. All AF3 models generated during the study have been deposited to ModelArchive repository (modelarchive.org).

### CDK5 kinase activity assay

The ADP-Glo assay (Promega) was used to study the kinase activity of CDK5 and CDK5-cyclin B1. Briefly, the kinase reaction was initiated by adding 5 μL of 7x ATP stock to 30 μL of a mixture containing CDK5 or CDK5-cyclin B1 and 50 μM model peptide substrate (KHHKSPKHA, purchased from Genscript). The final concentration of ATP in the reaction was 100 μM, and varied concentrations of the enzyme were used. The reaction was incubated for 2 hours at room temperature. The kinase activity was measured according to the manufacturer’s instructions with minor modifications. The ATPase reaction was stopped by adding and incubating with 30 μL of ADP-Glo reagent. Finally, 60 μL of kinase detection reagent was added to the reaction and incubated for an additional hour at room temperature before measuring the luminescence to quantify ATPase activity on a CLARIOstar microplate reader (BMG Labtech). All data were background-subtracted using buffer-only wells. Three independent experiments were performed in technical duplicates. Computed means and standard deviations were imported into Prism9 (GraphPad) and plotted as bar graphs.

### Mammalian Cell culture

All cells were cultured at 37 °C under a 5% CO2 atmosphere with 100% humidity. Telomerase-immortalized (hTERT) RPE-1 cells were cultured in DMEM-F12 medium (Thermo Fisher Scientific, 11320-033) containing 10% FBS (Thermo Fisher Scientific, A5256701) and 1% penicillin–streptomycin (Invitrogen, 15140-122). HeLa cells were cultured in DMEM with high glucose and pyruvate (Thermo Fisher Scientific, 11995-065) supplemented with 10% (v/v) FBS and 1% penicillin-streptomycin.

### Plasmids for cell-based assays

The CDK5, CDK1 and CDK2 glycine variants were generated using site-directed mutagenesis of the parental WT pOZ FLAG-HA plasmids by PCR using following primers: For CDK5-G152F (G-F)-forward, 5’-TTTATTCCCGTCCGCTGTTACTCAGCTGAGGTGGTCA CACTG-3’; reverse, 5’-CGGACGGGAATAAAAAAGGCTCGAGCCAGGCCAAAATCAGCCAATTTCAGCTCC-3’. For CDK5-G152W (G-W)-forward, 5’-TGGATTCCCGTCCGCTGTTACTCAGCTGAGGTGGTCACACTG-3’; reverse, 5’-CGGACGGGAATCCAAAAGGCTCGAGCCAGG CCAAAATCAGCCAATTTCAGCTCC-3’. For CDK1-G154F (G-F)-forward, 5’-TTTATCCCCA TCCGTGTGTACACCCACGAGGTGGTC-3’; reverse, 5’-CGGATGGGGATAAAGAAAGCAC GAGCCAGTCCGAAGTCAGCCAGCTTGATGG-3’. For CDK1-G154W (G-W)-forward, 5’-TGGATCCCCATCCGTGTGTACACCCACGAGGTGGTC-3’; reverse, 5’-CGGATGGGGATC CAGAAAGCACGAGCCAGTCCGAAGTCAGCCAGCTTGATGG-3’. For CDK2-G153F (G-F)-forward, 5’-TTTGTCCCTGTTCGTACTTACACCCATGAGGTGGTGACCCTGTG-3’; reverse, 5’-CGAACAGGGACAAAAAAAGCTCTGGCTAGTCCAAAGTCTGCTAGCTTGATGGCCCCCTC-3’. For CDK2-G153W (G-W)-forward, 5’-TGGGTCCCTGTTCGTACTTACACCCATGAGGTGGTGACCCTGTG-3’; reverse, 5’-CGAACAGGGACCCAAAAAGCTCTGGCTAGTCCAAAG TCTGCTAGCTTGATGGCCCCCTC-3’.

### Cell modification and related reagents

The generation of stable cell lines (CDK1/2/5 variant overexpressing lines) in this manuscript are described in detail below. RPE-1 cells stably expressing the pOZ Flag-HA-CDK1/2/5 variants (WT, G-F and G-W) were created by retroviral production and transduction. Retroviral production was performed using HEK293T Phoenix cells, which were co-transfected with pUMVC3-gag-pol, pCMV-VSVG/pCGEnv (Addgene, 8454), and pOZ-Flag-HA-CDK1/2/5 variant retroviral plasmids using Lipofectamine 2000 (Thermo Fisher Scientific, 11668019) according to the manufacturer’s instructions. Viral particles were collected 48h after transfection and filtered through a 0.45 μM membrane syringe filter (Thermo Fisher Scientific, 723-2545). Filtered viral supernatant was applied to RPE-1 cells in the presence of 8μg/ml polybrene (Sigma-Aldrich, 107689). As the pOZ is a retroviral vector that contains a bicistronic transcriptional unit that allows expression of the gene of interest and the selection marker (interleukin-2 receptor chain or IL-2Ra), transduced RPE-1 cells were selected by Dynabeads CD25, or anti-IL-2Rα antibody-conjugated magnetic beads (Thermo Fisher Scientific, 11157D).

### Cell cycle synchronization

For synchronization to mitotic prometaphase, cells were treated with nocodazole (Sigma Aldrich, M1404) dissolved in DMSO at the final concentration of 100 ng/ml for 6 hours. Cells were then collected by mitotic shake-off. For synchronization to S phase, cells were treated with 2 mM thymidine (Millipore Sigma, 6060-5GM) for 17 hours, washed with three times 1xPBS, and incubated in drug-free medium for 9 hours. Then, cells were incubated for a further 17 hours with 2 mM thymidine. Cells were released from double-thymidine by PBS washing and finally collected after 3 hours incubation in drug-free medium.

### Protein extraction, immunoblotting, and immunoprecipitation

Protein extracts were prepared in NETN buffer (50 mM Tris pH 7.5, 1 mM EDTA, 0.5% NP-40, 5% glycerol, 150 mM NaCl, cOmplete EDTA-free protease inhibitor cocktail (Sigma-Aldrich, 1183617001) and PhosSTOP phosphatase inhibitor cocktail (Sigma-Aldrich, 04906837001)). In brief, cells were collected and washed three times in ice-cold PBS before resuspending in lysis buffer. After incubation at 4 °C for 40 minutes with 10-second vortex every 10 minutes, the cell lysate was clarified by centrifuging at 13,000 rpm for 10 minutes at 4 °C. The protein concentration in clarified lysate was determined using the Pierce BCA Protein Assay Kit (Thermo Fisher Scientific, 23225) with reference to a standard curve generated with BSA.

Extracts were mixed with 4x SDS loading buffer (200 mM Tris-HCl pH 6.8, 8% (w/v) SDS, 0.1% (w/v) bromophenol blue, 40% (v/v) glycerol, and 400 mM 2-Mercaptoethanol) and heated at 95 °C for 10 minutes. Denatured extracts were resolved on precast NuPAGE 4-12%, 1.5 mm Bis-Tris polyacrylamide gels (Life Technologies, NP0335) and transferred to 0.2 μm nitrocellulose membranes (BioRad, 1620112). The membranes were blocked in 5% NFDM (Non Fat Dry Milk, Research Product International, M17200) in TBS-1% Tween-20 for 1 hour and incubated in primary antibodies (Vinculin, Santa Cruz Biotechnology sc-25336, 1: 200; CDK1/cdc2, Cell Signaling Technology 28439, 1:1000; CDK2, Abcam ab32147, 1:1000; CDK5, Abcam ab40773, 1:2000; Cyclin A2: rabbit Novus Biologicals NBP131330, 1:500; Cyclin B1: rabbit Cell Signaling Technology 4138, 1:1000; H3pS10: rabbit, Cell Signaling Technology 9701, 1:500; CDK1 pT14/pY15: rabbit, Thermo Fisher Scientific 44-686G, 1:1000; CDK5 pY15: rabbit, Cell Signaling Technology 19051, 1:1000) at 4 °C for 16–18 hours. The membranes were washed three times the next day with TBS-1% Tween-20 at room temperature with rocking. Membranes were probed with HRP-conjugated goat anti-mouse (Jackson Immuno Research, NC9832458) or goat anti-rabbit (Jackson Immuno Research, NC9736726) antibodies for 1 hour (1:5000). The Amersham ECL Western Blotting Detection Reagent Kit (GE Healthcare, RPN2106) was used to develop the blots. Blots were developed using Cytiva ImageQuant 800.

For CDK-cyclin binding investigation, synchronized cells were collected respectively and lysed as described above. Overexpressing CDK1/2/5 variants were immunoprecipitated by using Anti-FLAG (Sigma-Aldrich, A2220) agarose beads in 500-1000 μg protein lysate respectively in each pair investigation. The mixtures were incubated overnight with end-over-end rotation at 4 °C. The immunoprecipitants were then washed five times by NETN lysis buffer and mixed with 4x SDS loading buffer, followed by heating at 95 °C for 10 minutes. Eluates were examined by immunoblotting described as above.

## Statistical analyses

Statistical analyses were performed using Prism9 (GraphPad). All data are represented as mean ± SD, unless indicated otherwise.

## Extended Data Fig legends

**Extended Data Fig 1**. CDK5, CDK1 and CDK2 share extremely high sequence similarity with each other. A multiple sequence alignment (MSA) of CDK5, CDK1 and CDK2 was performed using Clustal Omega. Black-shaded letters indicate identical residue conservation, whereas grey-shaded letters show similar residues found at the same positions between aligned sequences. The critical regions and secondary structures are labelled on the MSA are based on the crystal structure of CDK5.

**Extended Data Fig 2**. Structural variations in CDK5-cyclin B1 complex models predicted by AF2 and AF3. a) The AF2-predicted model of CDK5-cyclin B1 complex with ipTM of 0.91. b) The AF3-predicted model of CDK5-cyclin B1 complex with ipTM of 0.87. Both AF2 and AF3 models are colored based on local model confidence indicated by pLDDT score for each amino acid and the color scale given at the bottom of the models. c) Superimposition of AF2 and AF3-predicted models of CDK5-cyclin B1 complex.

**Extended Data Fig 3**. CDK5-Cyclin B1 forms a complex in insect cells (Sf9 or HiFi). a) His-CDK5 co-purified with His-MBP-cyclin B1, when both proteins were co-expressed and purified using amylose resin to pull-down MBP-tagged cyclin B1 from HiFi or Sf9 cells. b) Reciprocally, His-cyclin B1 co-purified with His-MBP-CDK5, when both proteins were co-expressed and purified using amylose resin to pull-down MBP-tagged CDK5 from HiFi or SF9 cells.

**Extended Data Fig 4**. X-ray Crystal Structures of CDK5-Cyclin B1 Complexes: a) A 2.1 Å crystal structure of the CDK5-cyclin B1 complex in its apo form, with a 2Fo-Fc simulated annealing (SA) composite omit map shown near the CDK5 subunit (map is contoured at 1σ and carved at 2 Å near CDK5). b) A 2.5 Å crystal structure of the AMP-PNP-bound CDK5-cyclin B1 complex, with a 2Fo-Fc SA composite omit map shown near the cyclin B1 subunit (map is contoured at 1σ and carved at 2 Å near cyclin B1). c) A 2Fo-Fc SA composite omit map shown near critical regions in CDK5-cyclin B1-apo (C-helix and A-loop) and CDK5-cyclin B1-AMP-PNP (P-loop, hinge, and AMP-PNP). The map is contoured at 1σ and carved at 2 Å near the selected regions. In both structures, the P-loop and part (C157-L165) of the A-loop exhibit weak electron density/higher flexibility.

**Extended Data Fig 5**. The Alignment of hydrophobic catalytic-spine (C-spine) residues and AMP-PNP in the crystal structure of the active CDK5-cyclin B1-AMP-PNP complex. The C-spine is formed by two residues (V18 and A31) from β2 and β3 strands in N-ter lobe, AMP-PNP in the active site, three residues (L132-I134) from β8 strand right below the AMP-PNP, and additional three residues (L87, I191 and I195) from the C-ter lobe.

**Extended Data Fig 6**. Complex assembly and activation mechanisms are conserved among mitotic CDK-cyclin B1 complexes. a) Superposition of the CDK5-cyclin B1 crystal structure with the CDK2-cyclin A2 crystal structure (using cyclin as reference) reveals differences in CDK5 subunit disposition, A-loop stabilization, and active site configuration (distance between A48 and D84 is 23 Å in CDK2-cyclin A2 and 24 Å in the CDK5-cyclin B1). b) Superposition of the CDK5-cyclin B1 crystal structure with the AF2 model (using CDK as reference) reveals differences in cyclin subunit disposition, A-loop stabilization, and active site configuration (distance between A48 and D84 is 22 Å in AF2 and 24 Å in the crystal structure). c) Superposition of the CDK5-cyclin B1 crystal structure with the CDK2-cyclin A2 crystal structure (using CDK as reference) reveals differences in cyclin subunit disposition, A-loop stabilization, and active site configuration (distance between A48 and D84 is 23 Å in CDK2-cyclin A2 and 24 Å in the CDK5-cyclin B1).

**Extended Data Fig 7**. Sequence conservation of CDK5, CDK1 and CDK2 across different eukaryotic species. a) Logo plot showing consensus sequence of CDK5 generated from aligning 405 sequences of CDK5 across different species. b) Logo plot showing consensus sequence of CDK1 generated from aligning 494 sequences of CDK1 across different species. c) Logo plot showing consensus sequence of CDK2 generated from aligning 214 sequences of CDK2 across different species. Each letter represents the frequency of occurrence of a particular amino acid at that position. The conserved T14, Y15 and tip of the A-loop in CDK2, (and the corresponding positions CDK1 and CDK5) are highlighted with diamond symbols on CDK5, CDK1 and CDK2 consensus sequences.

**Extended Data Fig 8**. CDK5-cyclin B1 complex is regulated by post translation modification on CDK5 analogous to canonical CDK1-cyclin B1 complex as phosphorylation at T14, Y15 or both observed to affect active site formation. a) AF3 model of phos-T14-CDK5-cyclin B1-ATP complex colored based on the local confidence encoded by pLDDT score for each residue. b) AF3 model of phos-Y15-CDK5-cyclin B1-ATP complex colored based on the local confidence encoded by pLDDT score for each residue. c) AF3 model of phos-T14/phos-Y15-CDK5-cyclin B1-ATP complex colored based on the local confidence encoded by pLDDT score for each residue. d) AF3 model of phos-S159-CDK5-cyclin B1-ATP complex colored based on the local confidence encoded by pLDDT score for each residue.

**Extended Data Fig 9**. Inhibitory P-loop phosphorylation (pT14/pY15) on CDK1 and (pY15 on) CDK5 is diminished during mitosis. Endogenous CDK1 or CDK5 and other indicated proteins were immunoblotted from RPE1 and HeLa cells, collected at asynchronous population and mitosis in the cell cycle and stained by using indicated antibodies. Details of mitotic synchronization is described in detail in the Method section. Async., asynchronous, M., mitotic prometaphase.

## References

1 Nurse, P. A Long Twentieth Century of the Cell Cycle and Beyond. Cell 100, 71–78 (2000). 10.1016/s0092-8674(00)81684-0

2 Hochegger, H., Takeda, S. & Hunt, T. Cyclin-dependent kinases and cell-cycle transitions: does one fit all? Nature Reviews Molecular Cell Biology 9, 910–916 (2008). 10.1038/nrm2510

3 Malumbres, M. Cyclin-dependent kinases. Genome Biology 15 (2014). 10.1186/gb4184

4 Malumbres, M. & Barbacid, M. Mammalian cyclin-dependent kinases. Trends in Biochemical Sciences 30, 630–641 (2005). 10.1016/j.tibs.2005.09.005

5 Santamaría, D. et al. Cdk1 is sufficient to drive the mammalian cell cycle. Nature 448, 811–815 (2007). 10.1038/nature06046

6 Pao, P.-C. & Tsai, L.-H. Three decades of Cdk5. Journal of Biomedical Science 28 (2021). 10.1186/s12929-021-00774-y

7 Dhavan, R. & Tsai, L.-H. A decade of CDK5. Nature Reviews Molecular Cell Biology 2, 749–759 (2001). 10.1038/35096019

8 Zheng, X.-F. et al. A mitotic CDK5-PP4 phospho-signaling cascade primes 53BP1 for DNA repair in G1. Nature Communications 10 (2019). 10.1038/s41467-019-12084-x

9 Zheng, X.-F. et al. CDK5–cyclin B1 regulates mitotic fidelity. Nature 633, 932–940 (2024). 10.1038/s41586-024-07888-x

10 Brown, N. R., Noble, M. E. M., Endicott, J. A. & Johnson, L. N. The structural basis for specificity of substrate and recruitment peptides for cyclin-dependent kinases. Nature Cell Biology 1, 438–443 (1999). 10.1038/15674

11 Knighton, D. R. et al. Crystal Structure of the Catalytic Subunit of Cyclic Adenosine Monophosphate-Dependent Protein Kinase. Science 253, 407–414 (1991). 10.1126/science.1862342

12 Arter, C., Trask, L., Ward, S., Yeoh, S. & Bayliss, R. Structural features of the protein kinase domain and targeted binding by small-molecule inhibitors. Journal of Biological Chemistry 298 (2022). 10.1016/j.jbc.2022.102247

13 Kornev, A. P., Haste, N. M., Taylor, S. S. & Ten Eyck, L. F. Surface comparison of active and inactive protein kinases identifies a conserved activation mechanism. Proceedings of the National Academy of Sciences 103, 17783–17788 (2006). 10.1073/pnas.0607656103

14 Hu, J. et al. Kinase Regulation by Hydrophobic Spine Assembly in Cancer. Molecular and Cellular Biology 35, 264–276 (2023). 10.1128/mcb.00943-14

15 Tarricone, C. et al. Structure and Regulation of the CDK5-p25nck5a Complex. Molecular Cell 8, 657–669 (2001). 10.1016/s1097-2765(01)00343-4

16 Jeffrey, P. D. et al. Mechanism of CDK activation revealed by the structure of a cyclinA-CDK2 complex. Nature 376, 313–320 (1995). 10.1038/376313a0

17 Brown, N. R. et al. CDK1 structures reveal conserved and unique features of the essential cell cycle CDK. Nature Communications 6 (2015). 10.1038/ncomms7769

18 Jumper, J. et al. Highly accurate protein structure prediction with AlphaFold. Nature 596, 583–589 (2021). 10.1038/s41586-021-03819-2

19 Akdel, M. et al. A structural biology community assessment of AlphaFold2 applications. Nature Structural & Molecular Biology 29, 1056–1067 (2022). 10.1038/s41594-022-00849-w

20 Agarwal, V. & McShan, A. C. The power and pitfalls of AlphaFold2 for structure prediction beyond rigid globular proteins. Nature Chemical Biology 20, 950–959 (2024). 10.1038/s41589-024-01638-w

## References

1 Zheng, X.-F. et al. CDK5–cyclin B1 regulates mitotic fidelity. Nature 633, 932–940 (2024). 10.1038/s41586-024-07888-x

2 Gradia, S. D. et al. in DNA Repair Enzymes: Structure, Biophysics, and Mechanism Methods in Enzymology 1–26 (2017).

3 Kabsch, W. Xds. Acta Crystallographica Section D Biological Crystallography 66, 125–132 (2010). 10.1107/s0907444909047337

4 McCoy, A. J. et al. Phaser crystallographic software. Journal of Applied Crystallography 40, 658–674 (2007). 10.1107/s0021889807021206

5 Liebschner, D. et al. Macromolecular structure determination using X-rays, neutrons and electrons: recent developments in Phenix. Acta Crystallographica Section D Structural Biology 75, 861–877 (2019). 10.1107/s2059798319011471

6 Afonine, P. V. et al. Towards automated crystallographic structure refinement with phenix.refine. Acta Crystallographica Section D Biological Crystallography 68, 352–367 (2012). 10.1107/s0907444912001308

7 Emsley, P., Lohkamp, B., Scott, W. G. & Cowtan, K. Features and development of Coot. Acta Crystallographica Section D Biological Crystallography 66, 486–501 (2010). 10.1107/s0907444910007493

8 Meng, E. C. et al. UCSF ChimeraX: Tools for structure building and analysis. Protein Science 32 (2023). 10.1002/pro.4792

9 Bateman, A. et al. UniProt: the Universal Protein Knowledgebase in 2025. Nucleic Acids Research 53, D609–D617 (2025). 10.1093/nar/gkae1010

10 Bodenhofer, U., Bonatesta, E., Horejš-Kainrath, C. & Hochreiter, S. msa: an R package for multiple sequence alignment. Bioinformatics 31, 3997–3999 (2015). 10.1093/bioinformatics/btv494

11 Wagih, O. & Hancock, J. ggseqlogo: a versatile R package for drawing sequence logos. Bioinformatics 33, 3645–3647 (2017). 10.1093/bioinformatics/btx469

12 Abramson, J. et al. Accurate structure prediction of biomolecular interactions with AlphaFold 3. Nature 630, 493–500 (2024). 10.1038/s41586-024-07487-w

