## Extended Data Figures for "Human CDK5–cyclin B1 structures uncover a conserved mitotic CDK activation mechanism"

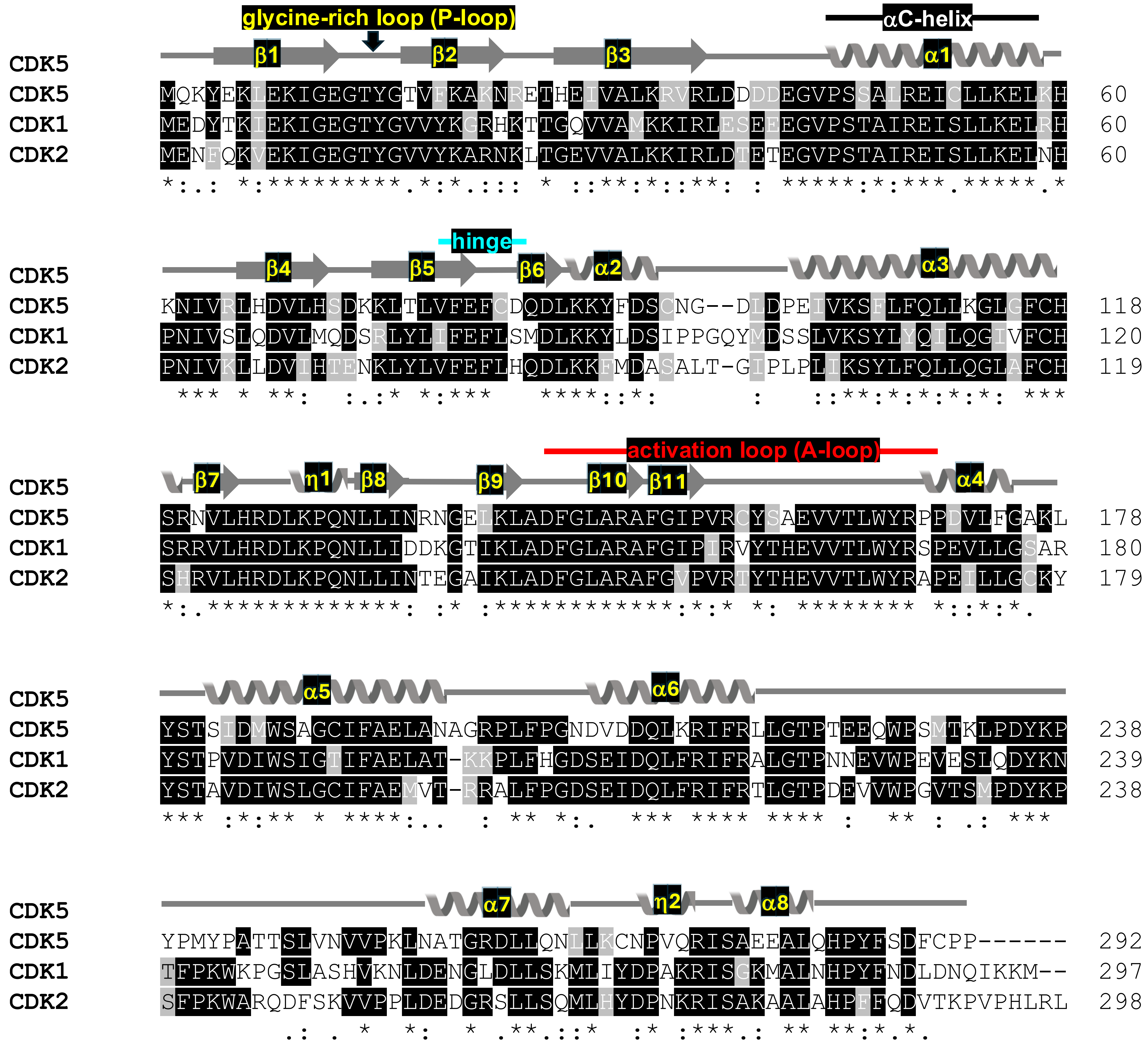


**Extended Data Fig 1.** CDK5, CDK1 and CDK2 share extremely high sequence similarity with each other. A multiple sequence alignment (MSA) of CDK5, CDK1 and CDK2 was performed using Clustal Omega. Black-shaded letters indicate identical residue conservation, whereas grey-shaded letters show similar residues found at the same positions between aligned sequences. The critical regions and secondary structures are labelled on the MSA are based on the crystal structure of CDK5.

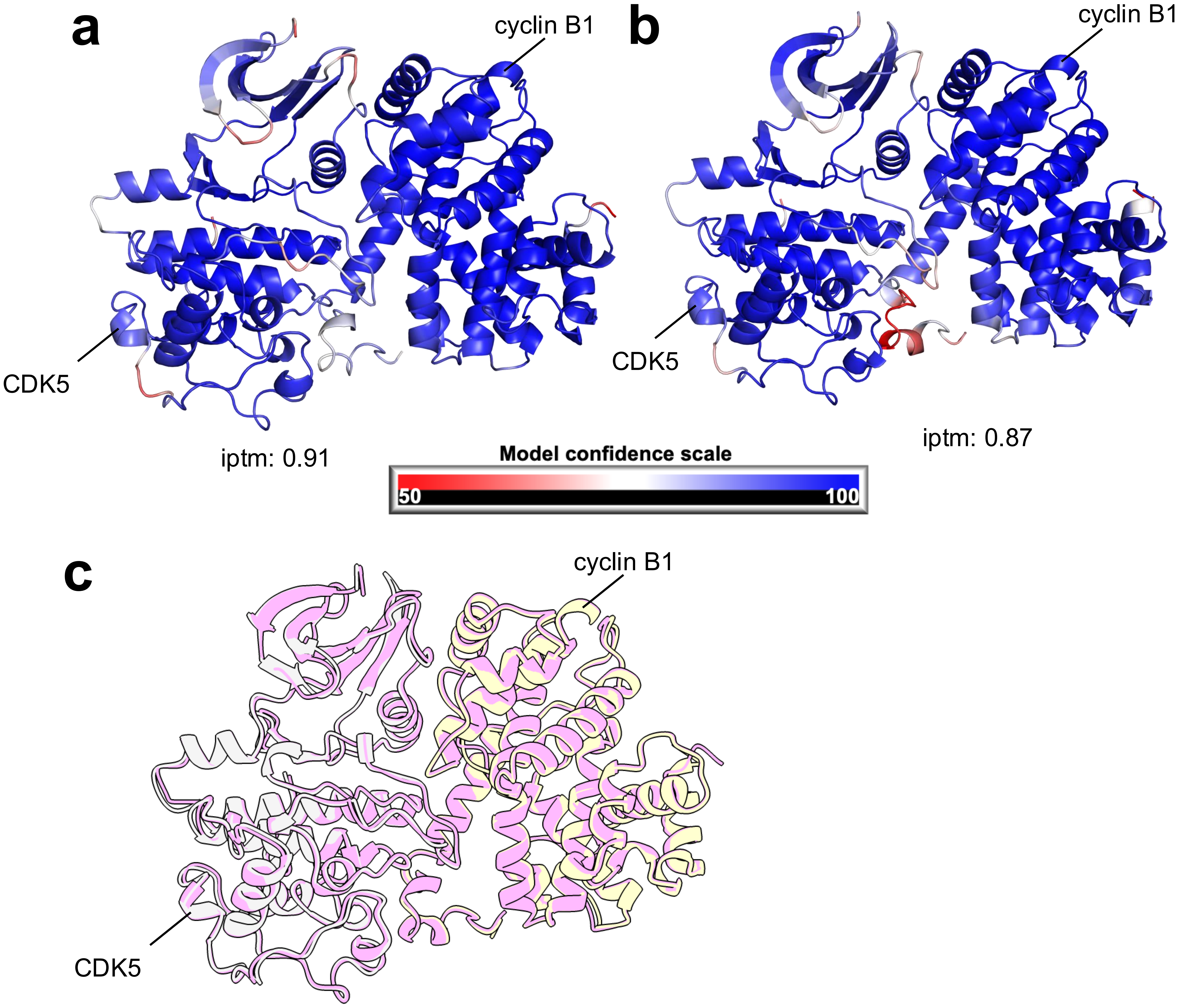


**Extended Data Fig 2.** Structural variations in CDK5-cyclin B1 complex models predicted by AF2 and AF3. a) The AF2-predicted model of CDK5-cyclin B1 complex with ipTM of 0.91. b) The AF3-predicted model of CDK5-cyclin B1 complex with ipTM of 0.87. Both AF2 and AF3 models are colored based on local model confidence indicated by pLDDT score for each amino acid and the color scale given at the bottom of the models. c) Superimposition of AF2 and AF3-predicted models of CDK5-cyclin B1 complex.

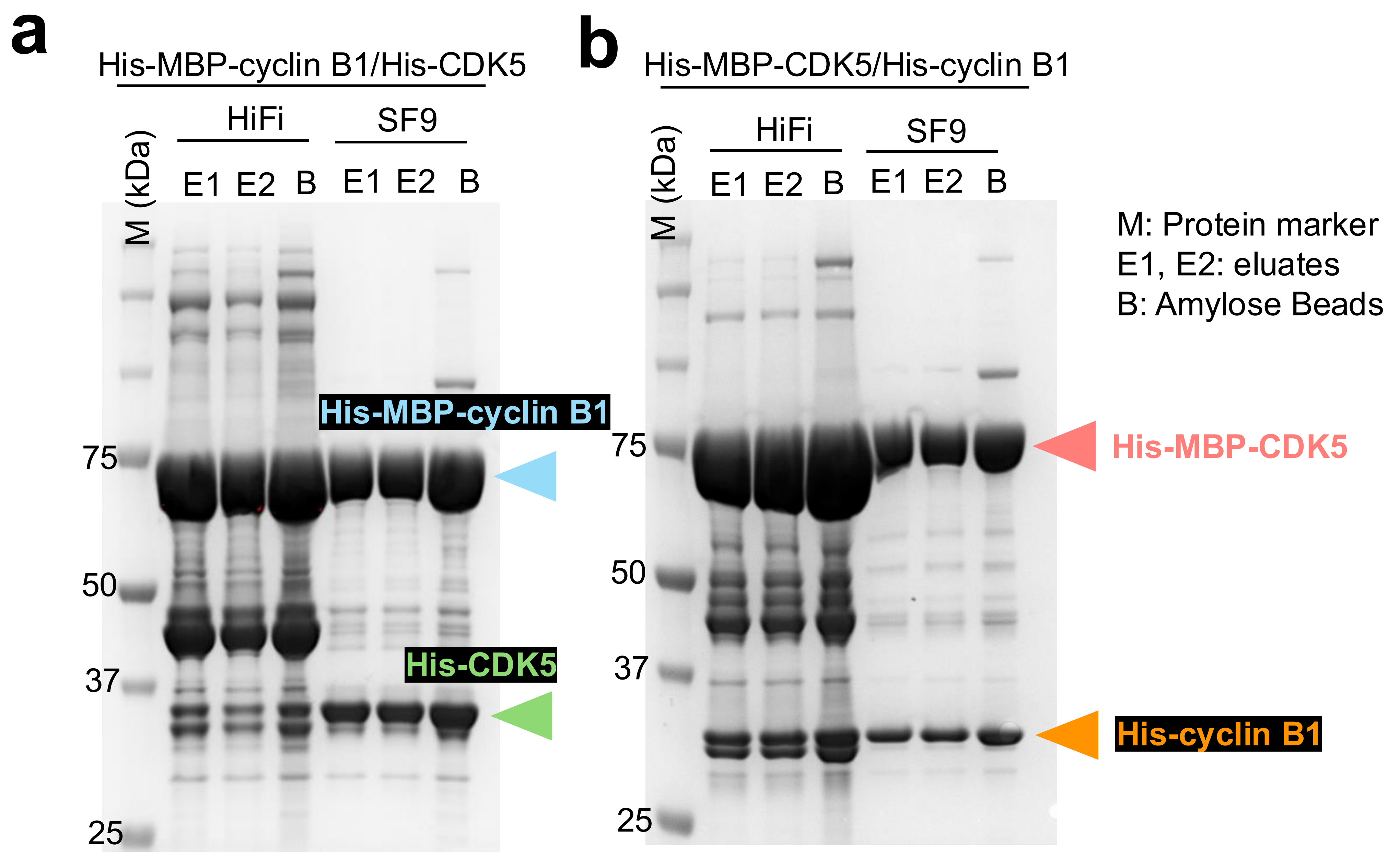


**Extended Data Fig 3.** CDK5-Cyclin B1 forms a complex in insect cells (Sf9 or HiFi). a) His-CDK5 co-purified with His-MBP-cyclin B1, when both proteins were co-expressed and purified using amylose resin to pull-down MBP-tagged cyclin B1 from HiFi or Sf9 cells. b) Reciprocally, His-cyclin B1 co-purified with His-MBP-CDK5, when both proteins were co-expressed and purified using amylose resin to pull-down MBP-tagged CDK5 from HiFi or SF9 cells.

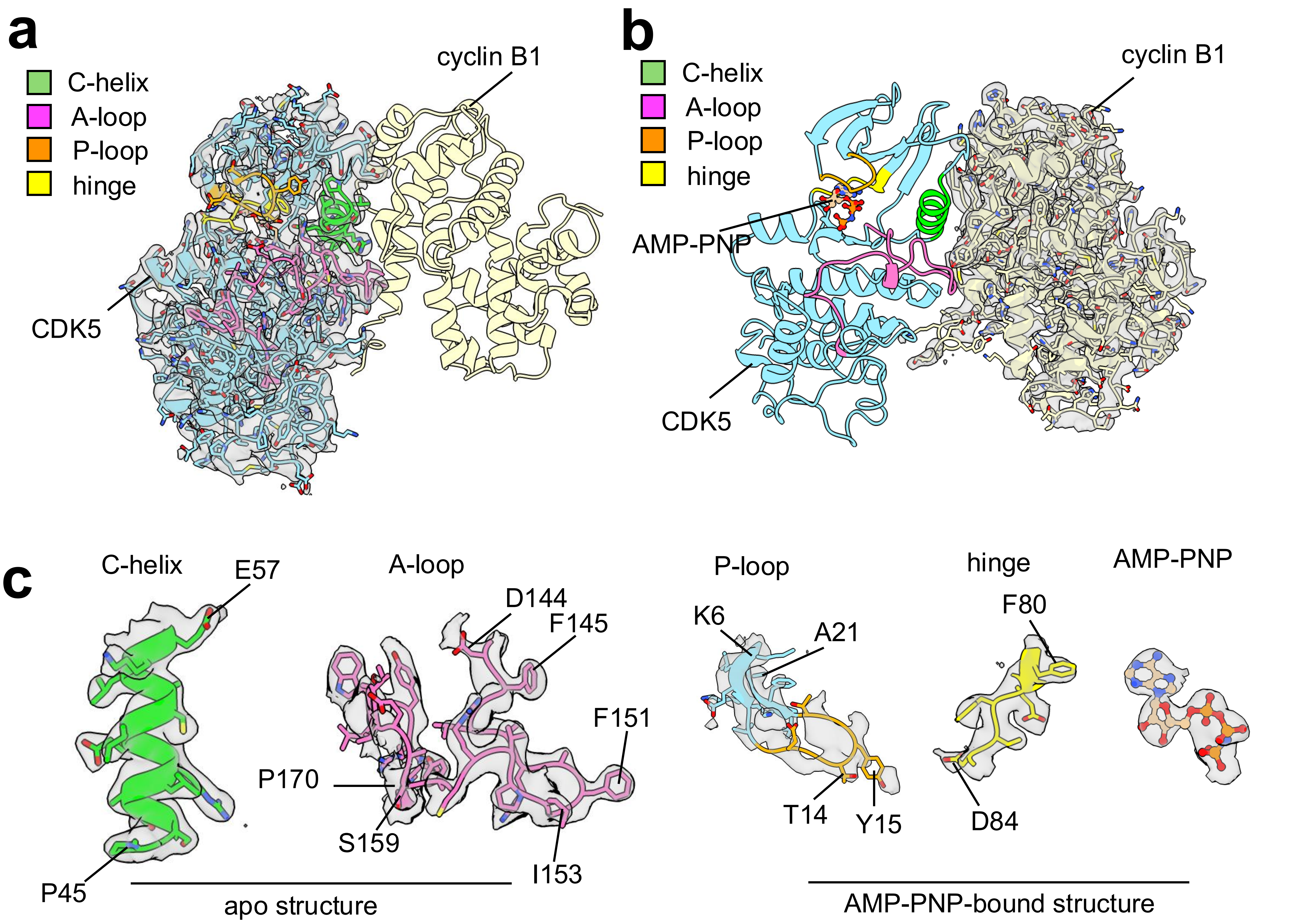


**Extended Data Fig 4.** X-ray Crystal Structures of CDK5-Cyclin B1 Complexes: a) A 2.1 Å crystal structure of the CDK5-cyclin B1 complex in its apo form, with a 2Fo-Fc simulated annealing (SA) composite omit map shown near the CDK5 subunit (map is contoured at 1σ and carved at 2 Å near CDK5). b) A 2.5 Å crystal structure of the AMP-PNP-bound CDK5-cyclin B1 complex, with a 2Fo-Fc SA composite omit map shown near the cyclin B1 subunit (map is contoured at 1σ and carved at 2 Å near cyclin B1). c) A 2Fo-Fc SA composite omit map shown near critical regions in CDK5-cyclin B1-apo (C-helix and A-loop) and CDK5-cyclin B1-AMP-PNP (P-loop, hinge, and AMP-PNP). The map is contoured at 1σ and carved at 2 Å near the selected regions. In both structures, the P-loop and part (C157-L165) of the A-loop exhibit weak electron density/higher flexibility.

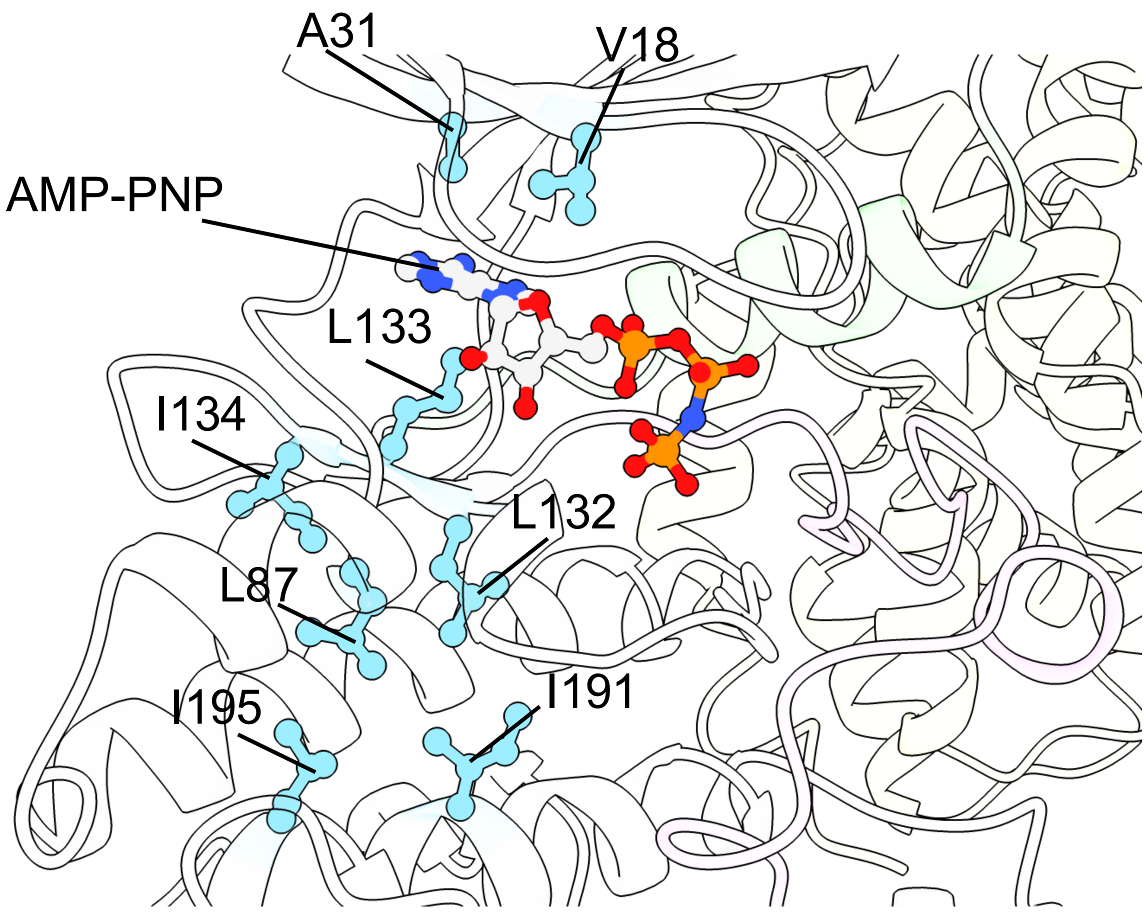


**Extended Data Fig 5.** The Alignment of hydrophobic catalytic-spine (C-spine) residues and AMP-PNP in the crystal structure of the active CDK5-cyclin B1-AMP-PNP complex. The C-spine is formed by two residues (V18 and A31) from β2 and β3 strands in N-ter lobe, AMP-PNP in the active site, three residues (L132-I134) from β8 strand right below the AMP-PNP, and additional three residues (L87, I191 and I195) from the C-ter lobe.

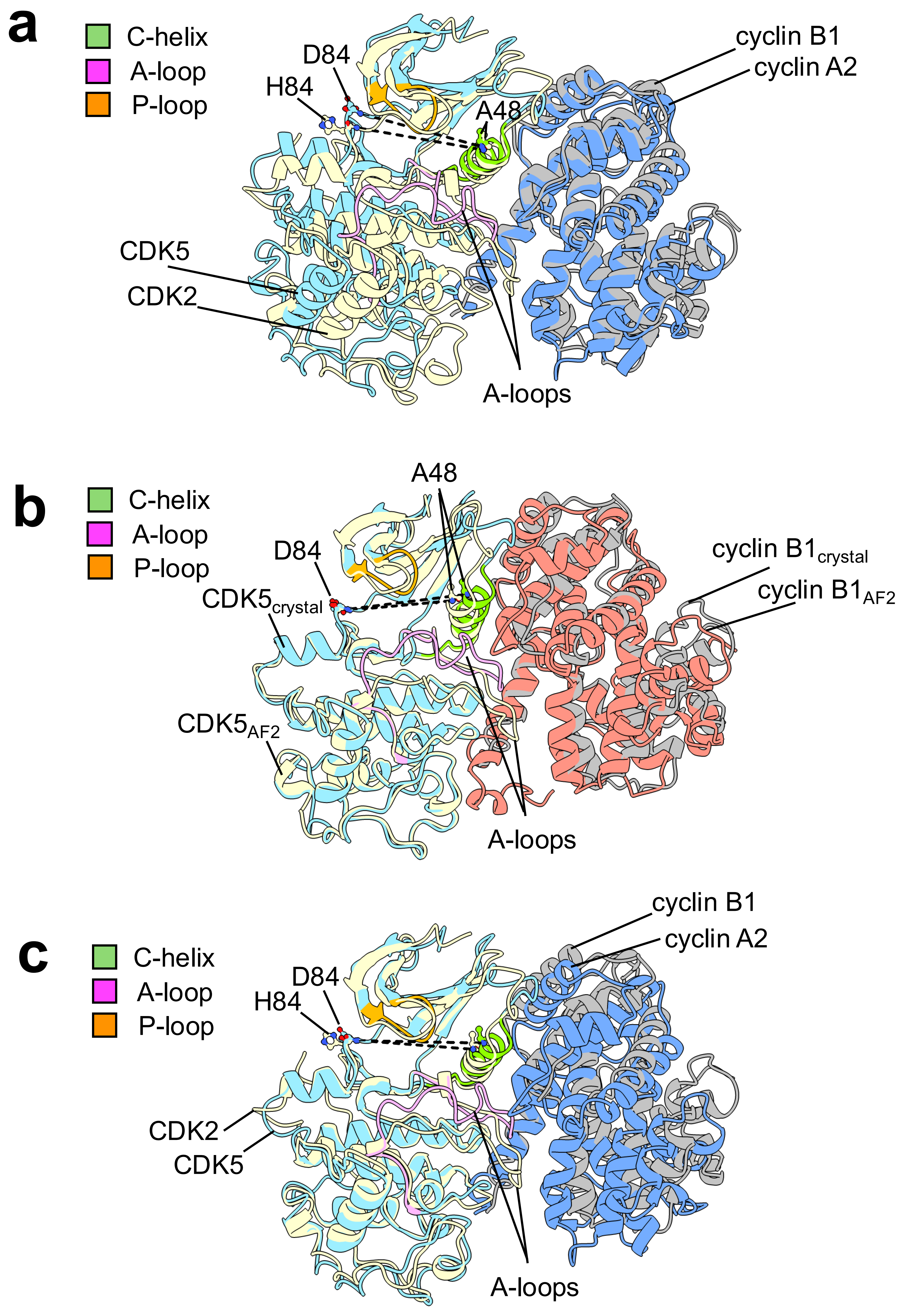


**Extended Data Fig 6.** Complex assembly and activation mechanisms are conserved among mitotic CDK-cyclin B1 complexes. a) Superposition of the CDK5-cyclin B1 crystal structure with the CDK2-cyclin A2 crystal structure (using cyclin as reference) reveals differences in CDK5 subunit disposition, A-loop stabilization, and active site configuration (distance between A48 and D84 is 23 Å in CDK2-cyclin A2 and 24 Å in the CDK5-cyclin B1). b) Superposition of the CDK5-cyclin B1 crystal structure with the AF2 model (using CDK as reference) reveals differences in cyclin subunit disposition, A-loop stabilization, and active site configuration (distance between A48 and D84 is 22 Å in AF2 and 24 Å in the crystal structure). c) Superposition of the CDK5-cyclin B1 crystal structure with the CDK2-cyclin A2 crystal structure (using CDK as reference) reveals differences in cyclin subunit disposition, A-loop stabilization, and active site configuration (distance between A48 and D84 is 23 Å in CDK2-cyclin A2 and 24 Å in the CDK5-cyclin B1).

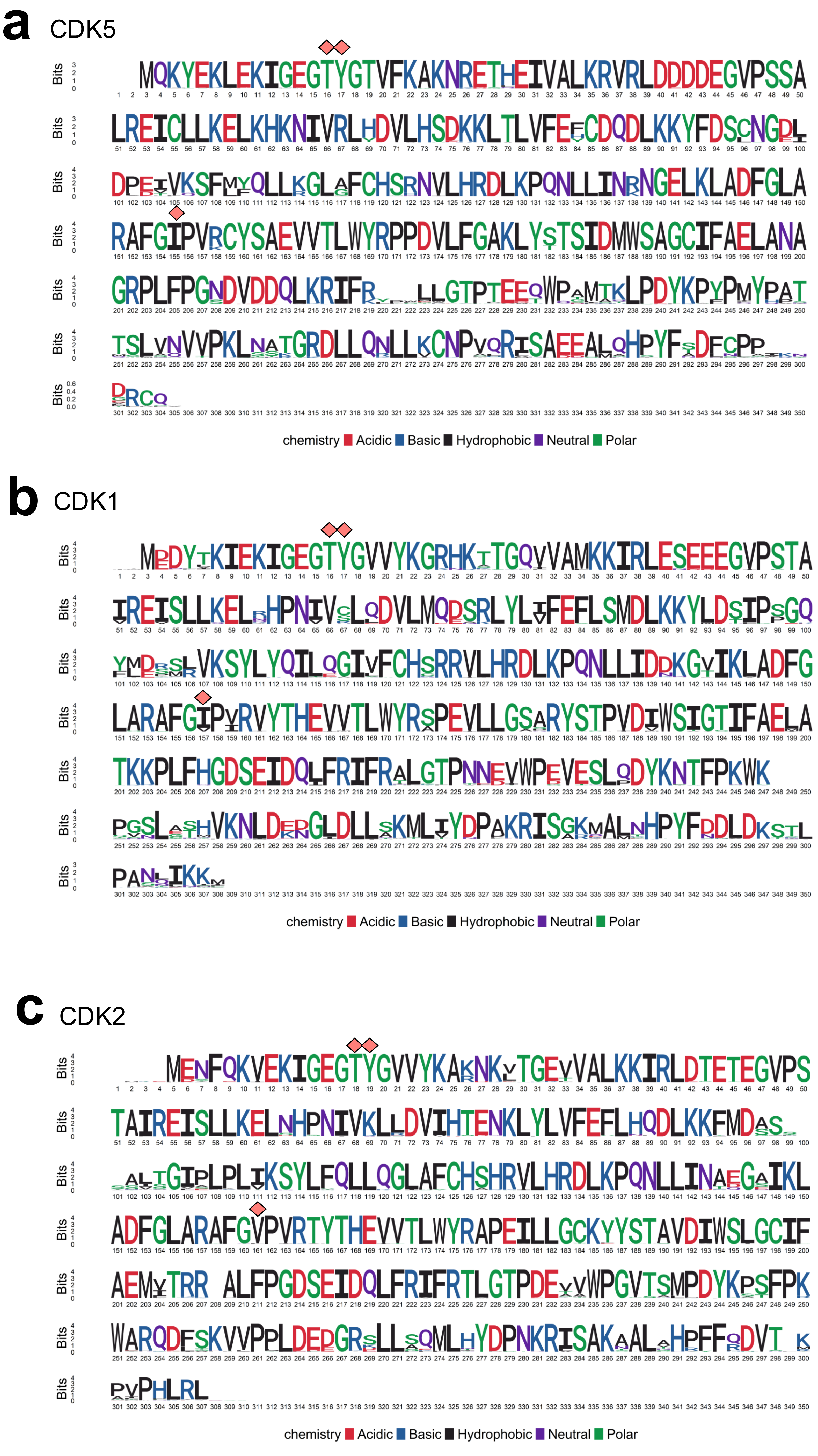


**Extended Data Fig 7.** Sequence conservation of CDK5, CDK1 and CDK2 across different eukaryotic species. a) Logo plot showing consensus sequence of CDK5 generated from aligning 405 sequences of CDK5 across different species. b) Logo plot showing consensus sequence of CDK1 generated from aligning 494 sequences of CDK1 across different species. c) Logo plot showing consensus sequence of CDK2 generated from aligning 214 sequences of CDK2 across different species. Each letter represents the frequency of occurrence of a particular amino acid at that position. The conserved T14, Y15 and tip of the A-loop in CDK2, (and the corresponding positions CDK1 and CDK5) are highlighted with diamond symbols on CDK5, CDK1 and CDK2 consensus sequences.

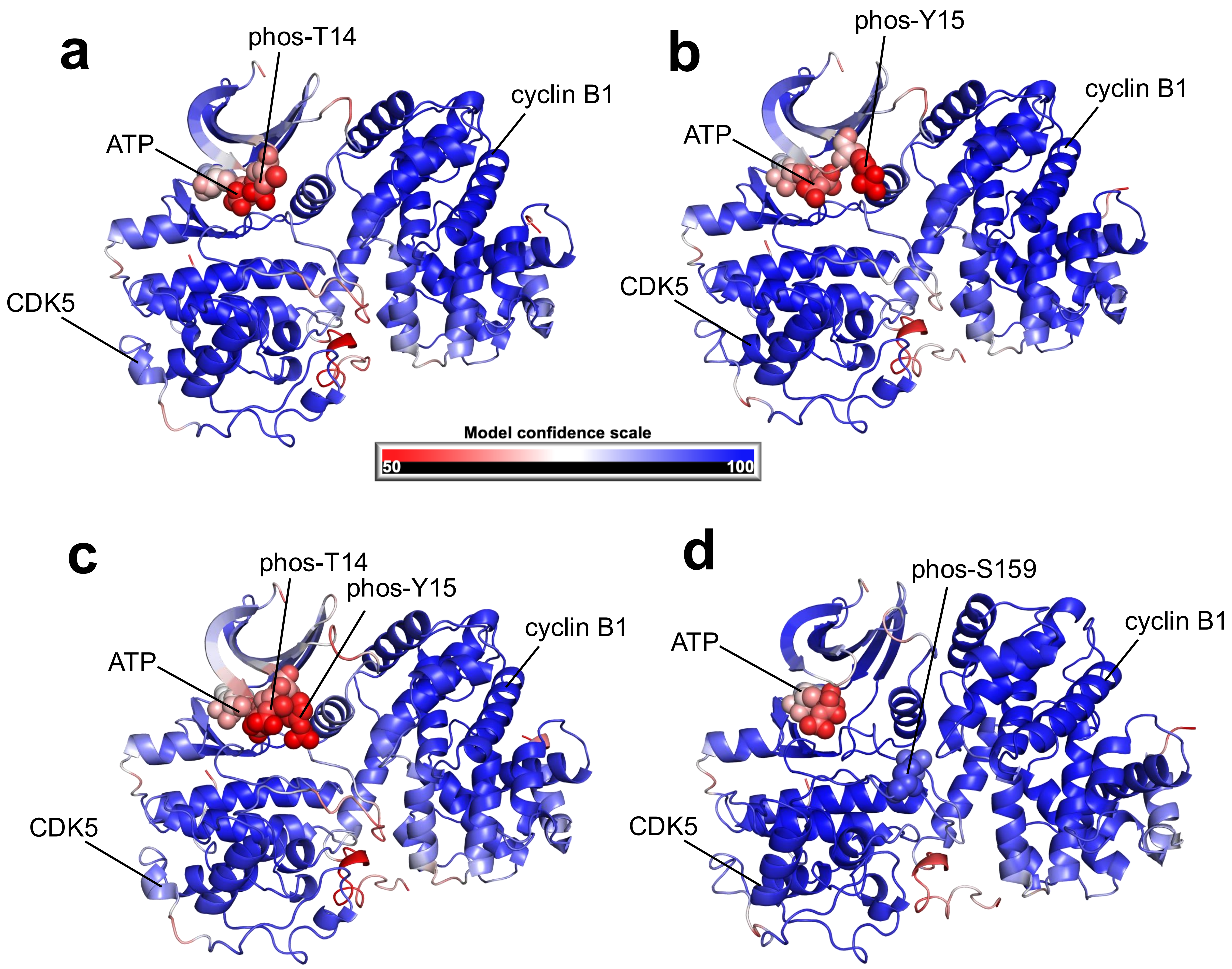


**Extended Data Fig 8.** CDK5-cyclin B1 complex is regulated by post translation modification on CDK5 analogous to canonical CDK1-cyclin B1 complex as phosphorylation at T14, Y15 or both observed to affect active site formation. a) AF3 model of phos-T14-CDK5-cyclin B1-ATP complex colored based on the local confidence encoded by pLDDT score for each residue. b) AF3 model of phos-Y15-CDK5-cyclin B1-ATP complex colored based on the local confidence encoded by pLDDT score for each residue. c) AF3 model of phos-T14/phos-Y15-CDK5-cyclin B1-ATP complex colored based on the local confidence encoded by pLDDT score for each residue. d) AF3 model of phos-S159-CDK5-cyclin B1-ATP complex colored based on the local confidence encoded by pLDDT score for each residue.

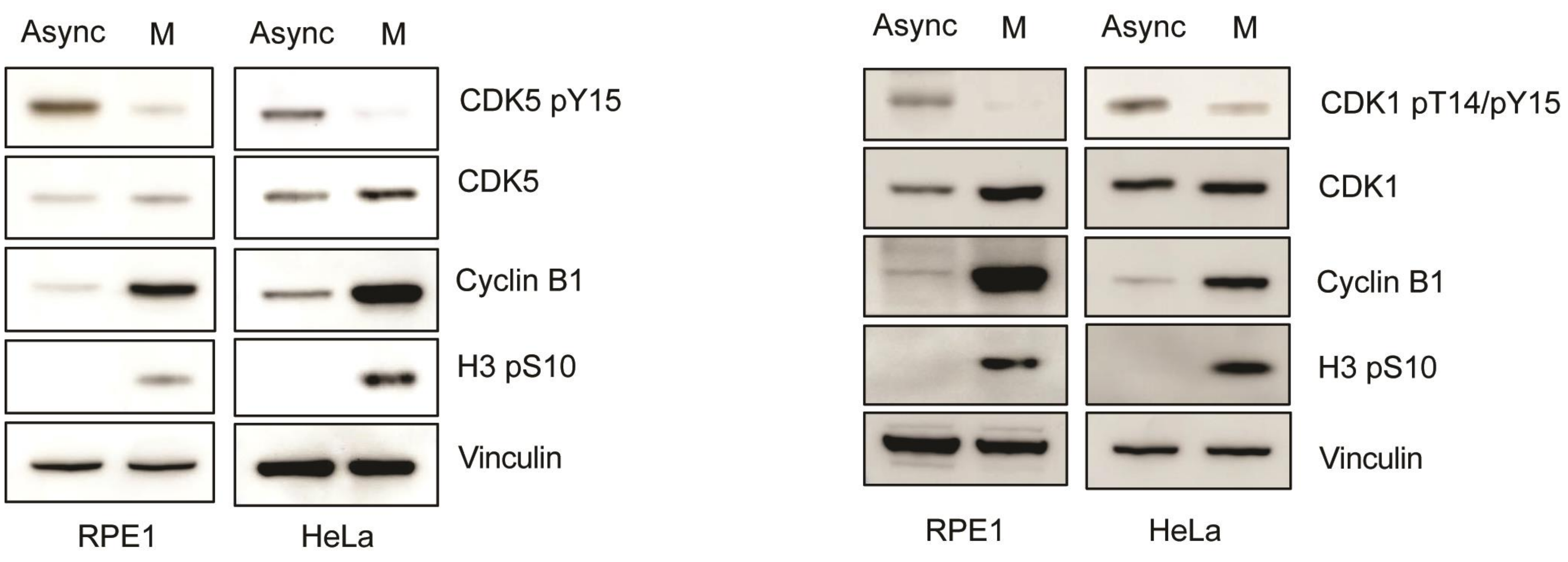


**Extended Data Fig 9.** Inhibitory P-loop phosphorylation (pT14/pY15) on CDK1 and (pY15 on) CDK5 is diminished during mitosis. Endogenous CDK1 or CDK5 and other indicated proteins were immunoblotted from RPE1 and HeLa cells, collected at asynchronous population and mitosis in the cell cycle and stained by using indicated antibodies. Details of mitotic synchronization is described in detail in the Method section. Async., asynchronous, M., mitotic prometaphase.
